# Economic Genome Assembly from Low Coverage Illumina and Nanopore Data

**DOI:** 10.1101/2020.02.07.939454

**Authors:** Thomas Gatter, Sarah von Löhneysen, Polina Drozdova, Tom Hartmann, Peter F. Stadler

## Abstract

We describe a new approach to assemble genomes from a combination of low-coverage short and long reads. `LazyBastard` starts from a bipartite overlap graph between long reads and restrictively filtered short-read unitigs, which are then reduced to a long-read overlap graph *G*. Edges are removed from *G* to obtain first a consistent orientation and then a DAG. Using heuristics based on properties of proper interval graphs, contigs are extracted as maximum weight paths. These are translated into genomic sequence only in the final step. A prototype implementation of `LazyBastard`, entirely written in python, not only yields significantly more accurate assemblies of the yeast and fruit fly genomes compared to state-of-the-art pipelines but also requires much less computational effort.

**Funding:** RSF / Helmholtz Association 18-44-06201; Deutsche Academische Austauschdienst, DFG STA 850/19-2 within SPP 1738; German Federal Ministery of Education an Research 031A538A, de.NBI-RBC

## Introduction

The assembly of genomic sequences from high throughput sequencing data has turned out to be a difficult computational problem in practice. Recent approaches combine cheap short-read data (typically using Illumina technology) with long reads produced by PacBio or Nanopore technologies. Although the short-read data are highly accurate and comparably cheap to produce, they are insufficient even at (very) high coverage due to repetitive elements. Long-read data, on the other hand, are comparably expensive and have much higher error rates. Similar to classical mate pair approaches, long reads can serve as “bridging elements” to resolve paths in classic (short read) assembly graphs [24]. Several assembly techniques have been developed recently for de novo assembly of large genomes from high-coverage (50× or greater) PacBio or Nanopore reads. Recent state-of-the-art methods employ a hybrid assembly strategy using Illumina reads to correct errors in the longer PacBio reads prior to assembly. For instance, the 32 Gb axolotl genome was produced in this manner [20]. Short reads then may serve for post-processing and correcting errors in a draft assembly [23]. The eel genome [9] was assembled using short reads as unique seeds that are phased and completed by Nanopore reads. While finishing this manuscript, `HASLR` [7] was published. Similar to our approach it uses unique contigs from short reads, however, goes on to define a backbone graph similar to the one defined by [9].

Here we describe an alternative approach to assembling genomes from a combination of long-read and short-read data. We strive to avoid the manipulation of global de Bruijn or string graphs as well as the direct all-against-all comparison of the error-prone long-read data and the difficult mapping of individual short reads against the long reads. As we shall see, this is not only possible but also adds the benefit of producing rather good assemblies with surprisingly low requirements on the coverage of both short and long reads.

## Strategy

Instead of a “total data” approach, we identify “anchors” that are nearly guaranteed to be correct and use an overall greedy-like workflow to obtain very large long-read contigs by “thinning-out” an initial overlap graph. The strategy of `LazyBastard` is outlined in Fig. 1. The key idea is to use a collection 𝒮 := {*s*_*i*_} of pre-assembled, high-quality sequences that are unique in the genome as “anchors” to determine overlaps among the long reads ℛ := {*r*_*j*_}. In practice, 𝒮 can be obtained by assembling Illumina data with fairly low coverage to the level of unitigs only. The total genomic coverage of 𝒮 only needs to be large enough to provide anchors between overlapping long reads, and it is rigorously filtered to be devoid of repetitive and highly similar sequences.

**Figure 1:**
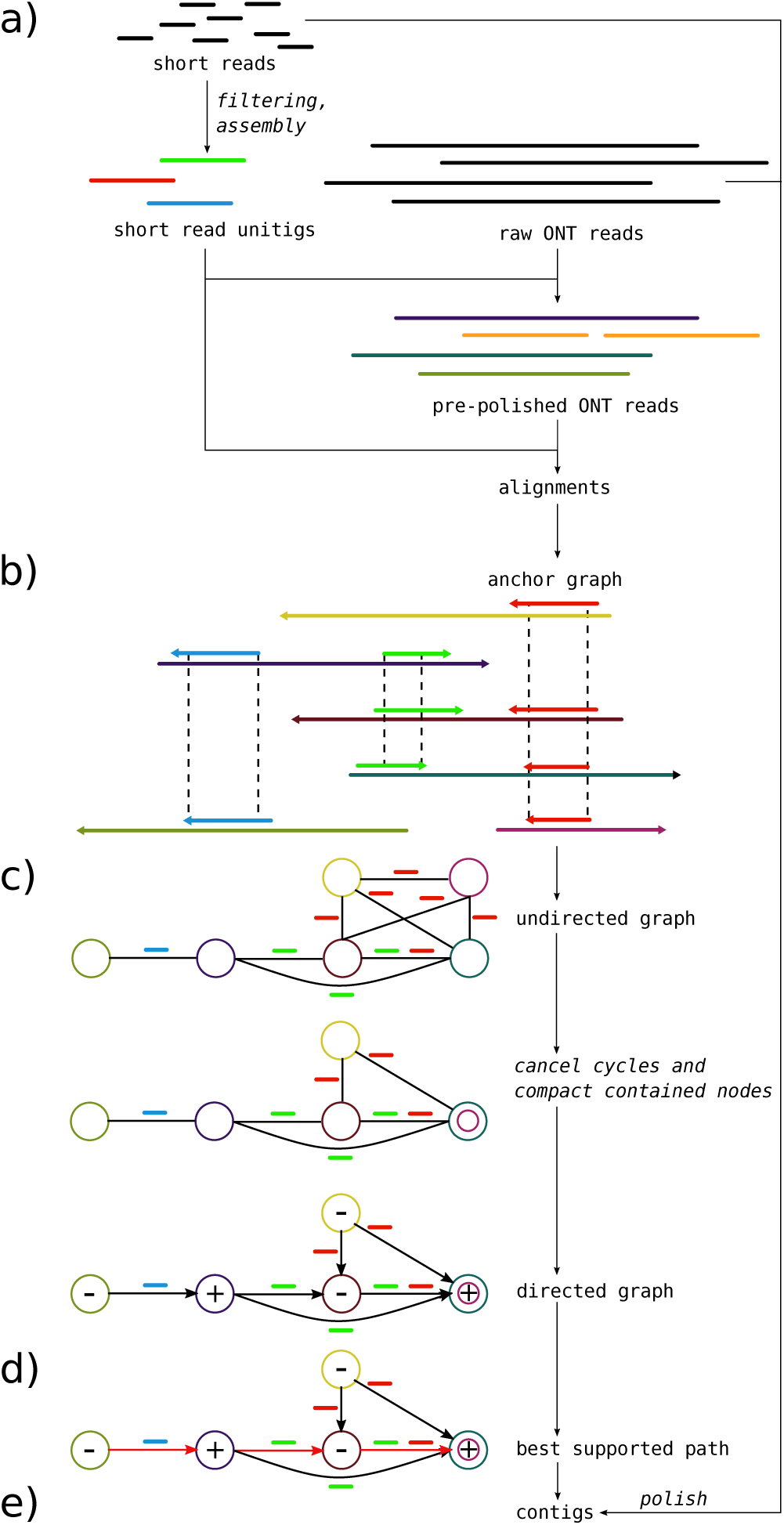
Overview of the `LazyBastard` assembly pipeline. a) Short Illumina reads are filtered to represent only near unique *k*-mers and subsequently assembled into unambiguous unitigs. Long Nanopore reads can be optionally scrubbed to include only regions consistent to at least one other read. For larger data sets scrubbing can be handled on subsets efficiently. Mapping unitigs against Nanopore reads yields unique “anchors” between them b). An undirected graph c) is created by adding Nanopore reads as nodes and edges between all pairs of reads sharing an “anchor”. Each edge is assigned a *relative orientation*, depending on whether the “anchor” maps in the same direction on both Nanopore reads. Cycles with a contradiction in orientation have to be removed before choosing a node at random and directing the graph based on its orientation. As Nanopore reads that are fully contained within another do not yield additional data, they can be collapsed. Contigs are extracted as maximally supported paths for each connected component d). Support in this context is defined by the number of consistent overlaps transitive to each edge. Final contigs e) can be optionally polished using established tools.

Mapping a short read *s* ∈ 𝒮 against the set ℛ of long reads implies (candidate) overlaps *r*_1_ − *r*_2_ between two long reads (as well as their relative orientation) whenever an *s* maps to both *r*_1_ and *r*_2_. Thus we obtain a directed overlap graph *G* of the long reads without an all-against-all comparison of the long reads. A series of linear-time filtering and reduction algorithms then prunes first the underlying undirected overlap graph and then the directed version of the reduced graph. Finally, contigs are extracted as best-supported path from its connected components. In the following section(s) we describe the individual steps in detail.

## Theory and Methods

### Preprocessing

A known complication of both PacBio and Nanopore technologies are chimeric reads formed by the artificial joining of disconnected parts of the genome [17] that thus may cause mis-assemblies [25]. Current methods dealing with this issue heavily rely on raw coverage [16] and hence are of little use for our goal of a low-coverage assembler. In addition, start- and end-regions of reads are known to be particularly error-prone. Nevertheless, we generally refrain from preprocessing the long reads at the outset and only consider problematic reads later at the level of the overlap graph.

Short-read (Illumina) data are preprocessed by adapter clipping and trimming. A set 𝒮 of high quality fragments is obtained from a restricted assembly of the short-read data. The conventional use case of assembly pipelines aims to find a minimal set of contigs in trade-off to both correctness and completeness. For our purposes, however, completeness is of little importance and fragmented contigs are not detrimental to our workflow, as long as their lengths stay above a statistical threshold. Instead, correctness and uniqueness are crucial. We therefore employ three filtering steps: (1) Using a *k*-mer profile, we remove all *k*-mers that are much more or much less abundant than the expected coverage since these are likely part of a repetitive sequence or contain sequencing errors. (2) In order to avoid ambiguities only branch-free paths are extracted from the assembly graph. Moreover, a minimal path length is required for secure anchors. The de Bruijn based assembler `ABySS` [21] allows to assemble up to unitig stage, implementing this goal. Since repeats in general lead to branch-points in the de Bruijn graph, repetitive sequences are strongly depleted in unitigs. While in theory, every such assembly requires a fine tuned *k*-mer size, we found overall results to be mostly invariant of this parameter. (3) Finally, the set ℛ of long reads is mapped against the unitig set. At present we use `minimap2` [15] for this purpose. Regions or whole unitigs significantly exceeding the expected coverage are removed from 𝒮 because they most likely are repetitive or at least belong to families of very similar sequences such as multi-gene families.

### Overlap Graph for Long Reads

As a result we obtain a set of *significant matches* 𝒱 := {(*s, r*) ∈ 𝒮 × ℛ 𝒮 *δ*(*s, r*) ≥ *δ*_*_} whose matching score *δ*(*s, r*) exceeds a user-defined threshold *δ*_*_. The *long-read overlap graph G* has the vertex set ℛ. Conceptually, two long reads overlap, i.e., there should be an undirected edge *r*_1_*r*_2_ ∈ *E*(*G*) if and only if there is an *s* ∈ 𝒮 such that (*s, r*_1_) ∈ 𝒱 and (*s, r*_2_) ∈ 𝒱. In practice, however, we employ a more restrictive procedure:

For distinct long reads *r*_1_, *r*_2_ ∈ ℛ with (*s, r*_1_), (*s, r*_2_) ∈ 𝒱 the sequence intervals on *s* that match intervals on *r*_1_ and *r*_2_ are denoted with [*i, j*] and [*k, l*], respectively. Without loss of generality let *i* ≤ *k*. The intersection [*i, j*] ∩ [*k, l*] is the interval [max{*i, k*}, min{*j, l*}] if *k* ≤ *j* and the empty interval otherwise. Note that if [*i, j*] ∩ [*k, l*] is not empty, then it corresponds to a direct match of *r*_1_ and *r*_2_. The expected bit score for the overlap is estimated as

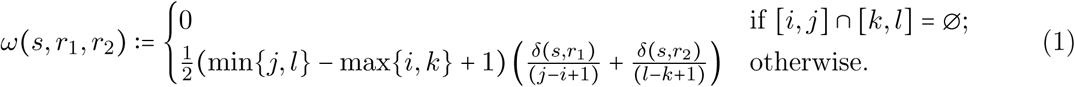

For a given edge *r*_1_*r*_2_ ∈ *E*(*G*) there may be multiple significant matches, mediated by a set of unitigs 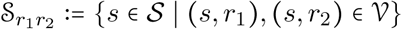. In ideal data they are all consistent with respect to orientation and co-linear location. In real data, however, this may not be the case.

For each significant match (*s, r*) ∈ 𝒱 we define the *relative orientation θ*(*s, r*) ∈ {+1, −1} of the reading directions of the short-read scaffold *s* relative to the long read *r*. The relative reading direction of the long reads (as suggested by *s*) is thus *θ*_*s*_(*r*_1_, *r*_2_) = *θ*(*s, r*_1_) · *θ*(*s, r*_1_). The position of a significant match (*s, r*) defined on the unitig *s* can be transferred to the long read *r* using a coordinate transformation *τ*_*r*_ (see Appendix). We write [*i, j*]_*r*_ := [*τ*_*r*_(*i*), *τ*_*r*_(*j*)] for the corresponding interval on *r*.

#### Definition 1.

*Two unitigs s, s*′ *in* 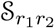 *are* consistent *if (i) θ*_*s*_(*r*_1_, *r*_2_) = *θ*_*s*′_ (*r*_1_, *r*_2_), *(ii) the relative order of* 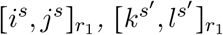 *on r*_1_ *and* 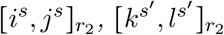 *on r*_2_ *is the same, and (iii) the difference between the distances of the intervals on r*_1_ *and r*_2_, *respectively, is sufficiently similar.*

We insert the edge *r*_1_*r*_2_ into *G* whenever the bit score Ω (*r*_1_, *r*_2_) := max_𝒮′_ ∑_*s*∈𝒮′_*ω*(*s, r*_1_, *r*_2_) computed over all maximal consistent sets of significant matches between *r*_1_ and *r*_2_ exceeds a user-defined threshold *δ*\*. Computing 𝒮′ is a chaining problem which is solved in quadratic time by dynamic programming [19]. The *signature θ*(*r*_1_, *r*_2_) of the edge *r*_1_*r*_2_ ∈ *E*(*G*) is the common value *θ*_*s*_(*r*_1_, *r*_2_) for all *s* ∈ 𝒮′.

For each edge *r*_1_*r*_2_ ∈ *E*(*G*) we determine 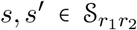 such that 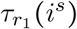 is the minimal and 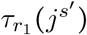 is the maximal coordinate of the matching intervals. The corresponding pair of coordinates, 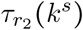 and 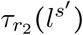, spans the matching intervals on (with upper and lower bound exchanged if *θ*(*r*_1_, *r*_2_) = −1). These intervals [*i, j*]_*r*_ specify the known overlapping regions between *r*_1_ and *r*_2_. If *θ*(*r*_1_, *r*_2_) = +1 then *r*_1_ extends *r*_2_ to the left if 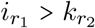 and to the right if 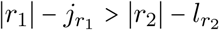. For *θ*(*r*_1_, *r*_2_) = −1 the corresponding conditions are 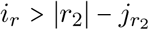 and 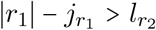, respectively. If *r*_2_ does not extend *r*_1_ to either side then *r*_2_ is completely contained in *r*_1_ and does not contribute to the assembly. We therefore contract the edge between *r*_1_ and *r*_2_. Otherwise, *r*_1_ extends *r*_2_ to the left and *r*_2_ extends *r*_1_ to the right, or *vice versa*. We record this direction as *r*_1_ → *r*_2_ or *r*_1_ ← *r*_2_, respectively.

The result of this construction is a long-read-overlap graph *G* whose vertices are the non-redundant long reads and whose edges *r*_1_*r*_2_ record (1) the relative orientation *θ*(*r*_1_, *r*_2_), (2) the bit score Ω (*r*_1_, *r*_2_), (3) the local direction of extension, and (4) the overlapping interval.

### Consistent Orientation of Long Reads

For perfect data it is possible to consistently determine the reading direction of each read relative to the genome from which it derives. This is not necessarily the case in real-life data. The relative orientation of two reads is implicitly determined by relative orientation of overlapping reads, i.e., by the signature *θ*(*r*_1_, *r*_2_) of the edge *r*_1_*r*_2_ ∈ *E*(*G*). To formalize this idea we consider a subset *D* ⊆ *E*(*G*) and define the *orientation* of *D* as 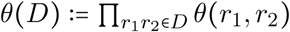. For a disjoint union of two edge sets we therefore have *θ*(*D* ⊍ *D*′) = *θ*(*D*′)*θ*(*D*), and more generally, the symmetric different *D* ⊕ *D*′ of two arbitrary edge sets *D* and *D*′ satisfies *θ*(*D* ⊕ *D*′) = *θ*(*D*)*θ*(*D*′) since the edges in *D* ∩ *D*′ appear twice in *θ*(*D*)*θ*(*D*′) and thus contribute a factor (±1)^2^ = 1.

#### Definition 2.

*Two vertices r*_1_*r*_2_ ∈ *V*(*G*) *are* orientable *if θ*(*P*) = *θ*(*P*′) *holds for any two paths P and P*′ *connecting r*_1_ *and r*_2_ *in G. We say that G is* orientable *if all pairs of vertices in G are orientable.*

#### Lemma 1.

*G is orientable if and only if every cycle C in G satisfies θ*(*C*) *=* 1.

*Proof.* Let *r, r*′ be two vertices of *G* and write 𝒞(*r, r*′) for the set of all cycles that contain *r* and *r*′. If *r* = *r*′ or 𝒞(*r, r*′) = Ø, then *r* and *r*′ are orientable by definition. Now assume *r* ≠ *r*′, 𝒞(*r, r*′) ≠ Ø, and consider a cycle *C* ∈ 𝒞(*r, r*′). Clearly, *C* can be split into two edge-disjoint path *C*_1_ and *C*_2_ both of which connect *r* and *r*′. If *r* and *r*′ are orientable, then *θ*(*C*_1_) = *θ*(*C*_2_) and thus *θ*_1_(*C*) = 1. If *r* and *r*′ are not orientable, then there is a pair of path *P*_1_ and *P*_2_ connecting *r* and *r*′ such that *θ*(*P*_1_) = −*θ*(*P*_2_). Since 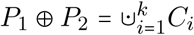 is an edge-disjoint union of cycles *C*_*i*_ we have 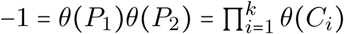 and thus there is least one cycle *C*_*i*_ with *θ*(*C*_*i*_) = −1 in *G.*

□

The practical importance of Lemma 1 is the implication that only a small set of cycles needs to be considered since every graph *G* with *c* connected components has a cycle basis comprising |*E*| − |*V*| − *c* cycles. Particular cycles bases, known as *Kirchhoff bases*, are obtained from a spanning tree *T* of *G* as the set ℬ of cycles *C*_*e*_ consisting of the edge *e* ∈ *E* \ *T* and the unique path *T* in connecting the endpoints of *e*. Every cycle *C* of *G* can then be written as *C* = ⊕_*e*∈*C*\*T*_ *C*_*e*_, see e.g. [11].

#### Theorem 1.

*Let* ℬ *be a cycle basis of G. The graph G is orientable if and only if θ*(*C*) = 1 *for all C* ∈ **B**.

*Proof.* The theorem follows from Lemma 1 and the fact that every cycle *C* in *G* can be written as an ⊕-sum of basis cycles, i.e., *θ*(*C*) = 1 for every cycle in *C* if and only if *θ*(*C*′) = 1 for every basis cycle *C*′ ∈ ℬ.

□

Theorem 1 suggests the following, conservative heuristic extracting an orientable subgraph from *G*:

1. Construct a maximum weight spanning tree *T*_*G*_ of *G* by using the Ω-scores as edge weights. Tree *T*_*G*_ can easily be obtained using, e.g., Kruskal’s algorithm [14].
2. Construct a Kirchhoff cycle basis ℬ from *T*_*G*_.
3. For every cycle *C* ∈ ℬ, check whether *θ*(*C*) = −1. If so, find the Ω-minimum weighted edge *ê* ∈ *C* and remove it from *E*(*G*) and (possibly) from *T*_*G*_ if *ê* ∈ *E*(*T*_*G*_). Observe that if *ê* ∉ *E*(*T*_*G*_), then *T*_*G*_ stays unchanged. If *ê* ∈ *E*(*T*_*G*_), then the removal of *ê* splits *T*_*G*_ into two connected components. We restrict *G* to the connected components of *T*_*G*_.

This procedure yields a not necessarily connected subgraph *G*′ and a spanning forest *T*_*G*_ ∩ *E*(*G*′) for *G*′.

#### Lemma 2.

*Let G be an undirected graph and let G*″ *be a connected component of the residual graph produced G*′ *by the heuristic steps (1)-(3). Then (i) G*″ *is orientable and (ii) T*_*G*_ ∩ *E*(*G*″) *is an* Ω*-maximal spanning tree of G*″.

*Proof.* Removal of an edge *e* from a spanning tree *T* of *G* partitions *T* into two components with vertex sets *V*_1_ and *V*_2_. Let *G*_1_ = *G*[*V*_1_] and *G*_2_ = *G*[*V*_2_] be the corresponding induced subgraphs of *G*. The cut in induced by *e* is *E*(*G*),(*E*(*G*_1_) ∪ *E*(*G*_1_)). Clearly *T*_1_ = *T* ∩ *E*(*G*_1_) and *T*_2_ = *T* ∩ *E*(*G*_2_) are spanning trees of *G*_1_ and *G*_2_, respectively. The restrictions ℬ_1_ and ℬ_1_ of the Kirchhoff basis ℬ to cycles with non-tree edges *e* ∈ *E*(*G*_1_) or *e* ∈ *E*(*G*_2_) form a Kirchhoff basis of *G*_1_ and *G*_2_, respectively. Now consider 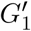 is obtained from *G*_1_ by removing a set of non-tree edges, then 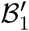 obtained from ℬ_1_ by removing the cycles with these non-tree edges is a cycle basis of 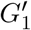 and *T*_1_ is still a spanning tree of *G*_1_. The Ω-weights of *T*_1_ = *T* ∩ *E*(*G*_1_) and *T*_2_ = *T* ∩ *E*(*G*_2_) must be maximal, since otherwise a heavier spanning tree *T* of *G* could be constructed by replacing *T*_1_ or *T*_2_ by a heavier spanning tree of *G*_1_ or *G*_2_. The arguments obviously extend to splitting *G*_*i*_ by cutting at an edge of *T*_*i*_. Since the heuristic removes all non-tree edges *e* with *θ*(*C*_*e*_) = −1, Theorem 1 implies that each component *G*″ is orientable. When removing *e* ∈ *T*, the corresponding cut edges are removed, the discussion above applies and thus *T* ∩ *E*(*G*″) is Ω-maximal.

□

From here on, we denote a connected component of *G*′ again by *G* and write *T*_*G*_ for its maximum Ω-weight spanning tree, which by Lemma 2 is just the restriction of the initial spanning tree to *G*. We continue by defining an orientation *φ* for the long reads. To this end, we pick an arbitrary *r*_*_ ∈ *V*(*G*) and set *φ* (*r*_*_) := +1. For each *r* ∈ *V*(*G*) we set 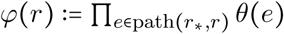, where path (*r*_*_, *r*) is the unique path connecting *r*_*_ and *r* in *T*_*G*_. We can now define an equivalent graph 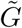 with the same vertices and edges as and orientations 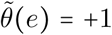 for *e* ∈ *T*_*G*_ and 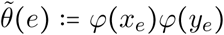 for all non-tree edges *e* = *x*_*e*_*y*_*e*_∉ *T*_*G*_. We note that the vertex orientations can be computed in *O*(|ℛ|) time along *T*_*G*_. Since 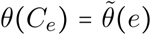 for every *C*_*e*_ ∈ ℬ, we can identify the non-orientable cycles in linear time.

### Golden Path

We now make use of the direction of the edges *r*_1_*r*_2_ defined by the mutual overhangs of the reads and write 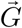 for the directed version of a connected component *G* of the residual graph *G*′ constructed above. For each edge *r*_1_*r*_2_ ∈ *E*(*G*) with *φ*(*r*_1_) = 1 we create the corresponding edge 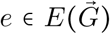 as follows:

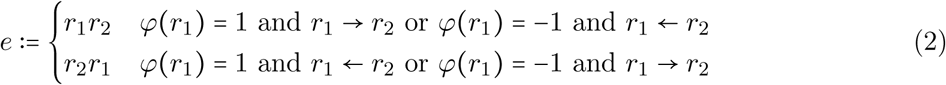

For *φ* = −1 the directions are reversed, i.e, *r*_1_ → *r*_2_ is replaced by *r*_2_ → *r*_1_ and *r*_1_ ← *r*_2_ by *r*_2_ ← *r*_1_.

In perfect data, 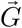 is a directed interval graph. Recall that we have contracted edges corresponding to nested reads (i.e., intervals). Therefore, 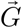 is a proper interval graph or indifference graph. Thus there is an ordering ≺ of the vertices (long reads) that satisfied the *umbrella property* [8]: *r*_1_ ≺ *r*_2_ ≺ *r*_3_ and 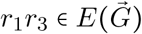 implies 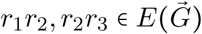. A “normal interval representation” and a linear order ≺ of the reads, can be computed in O(|ℛ|) time [18]. Again, we cannot use these results directly due to the noise in the original overlap graph.

First we observe that 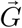 should be acyclic. Our processing so far, however, does not guarantee acyclicity since 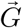 still may contain some spurious edges due to unrecognized repetitive elements. The obvious remedy is to remove a (weight-)minimal set of directed edges. This Feedback Arc Set problem, however, is NP complete, e.g., see [2] for a recent overview. We therefore resort to a heuristic that makes use of our expectations on the structure of 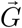: In general we expect multiple overlaps of correctly placed reads, i.e., *r* is expected to have several incoming edges from its predecessors and several outgoing edges exclusively to a small set of succeeding reads. In contrast, we expect incorrect edges to appear largely in isolation. This suggest to adapt Khan’s algorithm [10]. In its original version, it identifies a source *u*, i.e., a vertex with in-degree 0 is appended to the ordered vertex list *W* and all its out-edges are deleted. The algorithm stops with “fail” when no source can be found before the sorting is complete, indicating that a cycle has been encountered. In this case we consider as candidates the set *K* := (*V* \ *W*) ∩ *N*_+_(*W*) of not yet sorted out-neighbors of the vertices that are already sorted. Observe that *N*_+_(*W*) denotes the out-neighborhood of *W*. For each *u* ∈ *K* we distinguish incoming edges *xu* from *x* ∈ *W, x* ∈ *K*, and *x* ∈ *V* \ (*W* ∪ *K*). We distinguish two cases:

1. There is a *u* ∈ *K* without an in-edge *xu* from some other *x* ∈ *K*. Then we choose among these the *û* with the largest total Ω-weight incoming from *W* since this *û* overlaps with most of the previously sorted reads.
2. No such *u* exists, i.e., the candidate set *K* forms a strongly connected digraph. Then we choose the candidate *û* ∈ *K* with the largest difference of Ω-weights incoming from *W* and *K*.

In either case we remove the edges incoming from *V* \ *W* into *û* from 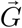 and proceed. We also modify the choice of the source such that we always pick the one with largest Ω-weight incoming from *W*. As a consequence, incomparable paths in 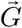 are sorted contiguously. The result of the modified Kahn algorithm clearly is an acyclic graph 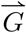.

For perfect data, 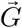 has a single source and a single sink vertex, corresponding to the left-most and right-most long reads *r*′and *r*″, respectively. Furthermore, every directed path connecting *r*′and *r*″ is a *golden path*, that is, a sequence of overlapping intervals that covers the entire chromosome. Even more stringently, every read *r* ≠ *r*′, *r*″ has at least one predecessor and at least one successor in 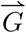. The acyclic graph 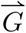 therefore has a unique topological sorting, i.e., its vertices are totally ordered. As before, we cannot expect that 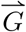 has these properties for real-life data.

A transitive reduction 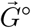 of directed graph 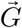 is a subgraph of 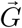 with as few edges as possible such that two vertices *x* and *y* are connected by a directed path in 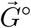 if and only if they are connected by a directed path in 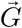. An acyclic digraphs 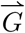 has a unique transitive reduction 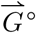 [1, Thm.1]. In the case of proper interval graphs, every redundant edge *uw* is part of a triangle due to the umbrella property. Consider a connected induced subgraph 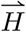 of the acyclic digraph 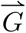 that is a proper interval graph. We observe that 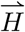 has a unique topological sorting and its triangle reduction 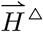, obtained by removing all edges *uw* for which there is a vertex *v* with 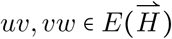, is a path. In fact, it is an induced path in the triangle reduction 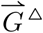 of 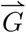. This suggests to identify maximal paths in 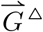 as contigs. Since the topological sorting along any such path is unique, it automatically identifies any redundant non-triangle edges along a path.

On imperfect data 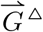 differs from a unique golden path by bubbles, tips, and crosslinks. Tips and bubbles predominantly are caused by edges that are missing e.g. due to mapping noise between reads that belong to a shared contig region. Hence, any path through a bubble or superbubble yields essentially the same assembly of the affected region and thus can be chosen arbitrarily, while tips may prematurely end the contig. Node-disjoint alternative paths within a (super-)bubble start and end in the neighborhood of the original path. Tips either originate or end in neighborhood of the chosen path.

Crosslinks represent connections between two proper contigs by spurious overlaps, caused e.g. by repetitive elements that have escaped filtering. As crosslinks can occur at any positions, a maximal path may not necessarily follow the correct connection and thus may introduce chimeras into the assembly. As a remedy we measure how well an edge *e* fits into a local region that forms an induced proper interval graph. Recall that the out-neighborhood of each vertex in a proper interval graph induces a transitive tournament. For real data, however, the subgraph 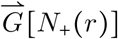 induced by the out-neighbors of *r* may in general violate this expectation. The problem of finding the maximum transitive tournament in an acyclic graph is NP-hard [4]. An approximation can be obtained, however, using the fact that a transitive tournament has a unique directed Hamiltonian path. Finding a longest path in a DAG only requires linear time. Thus candidates for transitive tournaments in 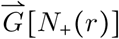 can be retrieved efficiently as the maximal path *P*_*rq*_ in 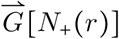 that connect *r* with an endpoint *q*, i.e., a vertex without an outgoing edge within 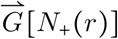. Clearly, it suffices to consider the maximum path problem in the much sparser 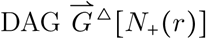. The induced subgraph 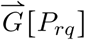 with the largest edge set 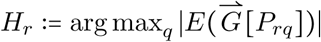 serves as approximation for the maximal transitive tournament and is used to define the *interval support* as

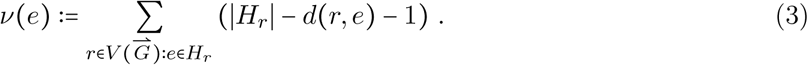

Here, *d*(*r, e*) is the minimal number of edges in a path from *r* to *e* in 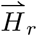. The interval support can be interpreted as the number of triangles that support *e* as lying within an induced proper interval graph. It suffices to compute *ν*(*e*) for 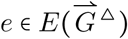. Empirically we observed that *ν*(*e*) is better than the weight Ω of the spanning tree edges in determining the best path. Taken together, we arrive at the following heuristic to iteratively extract meaningful paths:

i. Find the longest path **p** = *r*_1_,…, *r*_*n*_ in 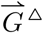 such that at every junction, we choose the incoming and outgoing edges *e* with maximal *ν*(*e*).
ii. Add the path **p** to the contig set if it is at least two nodes long and neither the in-neighborhood *N*_−_(*r*_1_) nor the out-neighborhood *N*_+_(*r*_1_) are marked as previously visited in 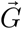. Otherwise, we have found a tip if one of *N*_−_(*r*_1_) or *N*_+_(*r*_1_) was visited before and a bubble if both were visited. Such paths are assumed to have arisen from more complex crosslinks and can be added to the contig set if they exceed a user-defined minimum length.
iii. The path **p** is marked visited in 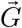 and all corresponding nodes and edges are deleted from 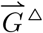.
iv. The procedure terminates when 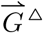 is empty.

As the result, we obtain a set of paths, each defining a contig.

### Consensus Sequence

Finally, we find the consensus sequence for each path **p**. This step is more complicated than usual due to the nature of our initial mappings. While we enforce compatible sets of unitigs for each pair of long reads, a shared unitig between edges does not necessarily imply the same genomic coordinate. (i) Unitigs can be long enough that we gain triples 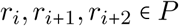 such that an 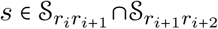 exists but *r*_*i*_ and *r*_*i*+2_ share no interval on *s*. Such triples can occur chained. (ii) Unitigs of genomic repeats may remain in the data. Such unitigs may introduce triples of edges *e*_*i*_ ≺ *e*_*j*_ ≺ *e*_*j*_ along the path *P*, where 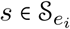 and 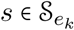 but 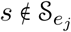, therefore creating disconnected occurrences of. (iii) Similarly, proximal repeats may cause inversions in the order of two unitigs 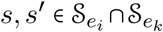 w.l.o.g *e*_*i*_ ≺ *e*_*j*_. This scenario cannot appear on neighboring edges, as the shared node has a unique order of *s* and *s*′. Hence, either *s* or *s*′ must be missing in an intermediary edge *e*_*l*_ due to the consistency constraints in the original graph, resulting in a situation as described in (ii). (iv) Finally, true matches of unitigs may be missing for some long reads due to alignment noise, which may also yield a situation as in (ii). To address (i), we collect all instances of a unitig in the path independent of its context. We create an undirected auxiliary graph *U*_*s*_ with a vertex set *V*(*U*_*s*_) := {*v* :=*e* | ∀*e* ∈ *E*(*P*) : *s* S_*e*_}. We add edges for all edge-pairs that share an overlap in *s*. Any clique in this graph then represents a set of edges that share a common interval in *s*. We assign each edge a unique cluster index 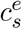, according to a minimal size clique decomposition. As finding a set of maximal cliques is NP-hard, we instead resort to a *O*(|*V*|/(log|*V*|)^2^) heuristic [3]. We address (ii-iv) with the help of a second index 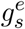, where 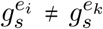 for two edges *e*_*i*_, *e*_*k*_ if and only if an edge *e*_*j*_ exists s.t. *e*_*i*_ ≺ *e*_*j*_ ≺ *e*_*j*_ and 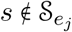.

Finally, we can now create a multigraph *M* consisting of vertex triples 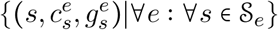. We add edges 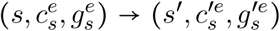 if and only if *s* ≺ *s*′on an edge *e* and no element *s*″ exists such that *s* ≺ *s*″ ≺ *s*′. The resulting graph is cycle free and thus uniquely defines the positions of all unitigs. Nodes represent the sequence of the common interval on the unitig *s* as attributed to the clique 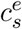. Edges represent the respective sequence of long reads between *s* and *s*′, or a negative offset value if unitigs overlap. We take an arbitrary node in *M* and set its interval as the reference point. Positions of all other nodes are progressively built up following a topological order in this graph. If multiple edges exist between two nodes in this process a random but fixed edge is chosen to estimate the distance between nodes. As now all sequence features are embedded in the same coordinate system, an arbitrary projection of the sequence is set as the reference contig, retaining unitigs were possible due to their higher sequence quality. At the same time, we can map the features of each long read to their respective position in this newly constructed reference. This information can be directly fed into consensus based error correction systems such as `racon` [23].

## Experimental Results

To demonstrate the feasibility of our assembly strategy we applied `LazyBastard` to publicly available datasets [5, 12, 22] for three well studied model organisms, baker’s yeast (*S. cerevisiae*, genome size 12 Mb), fruit fly (*D. melanogaster*, genome size 140 Mb) and human (*H. sapiens*, genome size 3GB). The data were downsampled to approximately 5× and 10× nanopore coverage for long reads, respectively, and Illumina coverage sufficient for short-read anchors. We compare results to the most widespread competing assembler `Canu` [13], as well as the recent competitor `HASLR` [7] implementing a similar concept. For comparison, we also provide the statistics for short-read only assemblies created with `ABySS` [21] on the same sets of reads used to create the “anchors”. Quality was assessed via alignment to a reference genome by the `QUAST` tool [6]; see Tbl. 1. `LazyBastard` produced consistently better results than `Canu`, increasing genomic coverage at a lower contig count. Due to our inclusion of accurate short-read unitigs, overall error counts are also significantly lower. Most notably, `Canu` was unable to properly operate at the 5× mark for both data sets. Only insignificant portions of yeast could be assembled, accounting for less than 15% of the genome. `Canu` completely failed for fruit fly, even after adapting settings to low coverage. Even at 5×, `LazyBastard` already significantly reduces the number of contigs compared to the respective short-read assemblies, while retaining a reasonably close percentage of genome coverage. At only 10× coverage for fruit fly, we were able to reduce the contig count 10-fold at better error rates. For human, `LazyBastard` manages at 39-fold decrease of the number contigs, albeit at a loss of greater 10% coverage. This difference appears to be a consequence of the high fragmentation of unitigs in the abundant repeat regions of the genome, rendering them too unreliable as anchors. Results are indeed in line with unitig coverage. While `HASLR` produced the fewest mis-assemblies, it creates significantly more and shorter contigs that cover a much smaller fraction of the genome. As a consequence it has the least favorable N50 values of all tools. For fruit fly at 10×, it results in four times as many contigs and covers 10% less of the genome, with a 12 times lower N50. While an improvement to `Canu`, it also struggles on datasets with low Nanopore coverage. We could not process our human test set with `HASLR`.

**Table 1:** Assessment of assembly qualities for `LazyBastard`, `Canu`, and short-read only assemblies for two model organisms. `LazyBastard` outperforms `Canu` in all categories, while significantly reducing contig counts compared to short-read only assemblies. While `HASLR` is more accurate, it covers significantly less of genomes at a higher contig count and drastically lower N50. Mismatches and InDels are given per 100kb. Accordingly, errors in `LazyBastard`’s unpolished output constitute <1% except for human. Column descriptions: X coverage of sequencing data, **compl**eteness of the assembly. **#ctg** number of contigs, **#MA** number of misassemblies (breakpoints relative to the reference assembly) **M**is**M**atches and **InDels** relative to the reference genomes. N50 of correctly assembled contigs (minimal length contig needed to cover 50% of the genome).

| Organism | X | Tool | compl.[%] | #ctg | #MA | MM | InDels | N50 |
| --- | --- | --- | --- | --- | --- | --- | --- | --- |
| yeast | ~5× | LazyB | 90.466 | 127 | 9 | 192.56 | 274.62 | 118843 |
|  |  | Canu | 14.245 | 115 | 5 | 361.47 | 2039.15 | - |
|  |  | SLR | 64.158 | 111 | 1 | 14.87 | 34.86 | 60316 |
|  | ~11× | LazyB | 97.632 | 33 | 15 | 193.73 | 300.20 | 505126 |
|  |  | Canu | 92.615 | 66 | 15 | 107.00 | 1343.37 | 247477 |
|  |  | SLR | 92.480 | 57 | 1 | 7.89 | 33.91 | 251119 |
|  | ~80× | Abyss | 95.247 | 283 | 0 | 9.13 | 1.90 | 90927 |
| fruit fly | ~5× | LazyB | 71.624 | 1879 | 68 | 446.19 | 492.43 | 64415 |
|  |  | Canu | - | - | - | - | - | - |
|  |  | SLR | 24.484 | 1407 | 10 | 31.07 | 58.96 | - |
|  | ~10× | LazyB | 80.111 | 596 | 99 | 433.37 | 486.28 | 454664 |
|  |  | Canu | 49.262 | 1411 | 275 | 494.66 | 1691.11 | - |
|  |  | SLR | 67.059 | 2463 | 45 | 43.83 | 84.89 | 36979 |
|  | ~45× | Abyss | 83.628 | 5811 | 123 | 6.20 | 8.31 | 67970 |
| human | ~10× | LazyB | 67.108 | 13210 | 2915 | 1177.59 | 1112.84 | 168170 |
|  |  | Canu | - | - | - | - | - | - |
|  | ~43× | Unitig | 69.422 | 4146090 | 252 | 93.07 | 13.65 | 338 |
|  | ~43× | Abyss | 84.180 | 510315 | 2669 | 98.53 | 25.03 | 7963 |

**Table 2:**
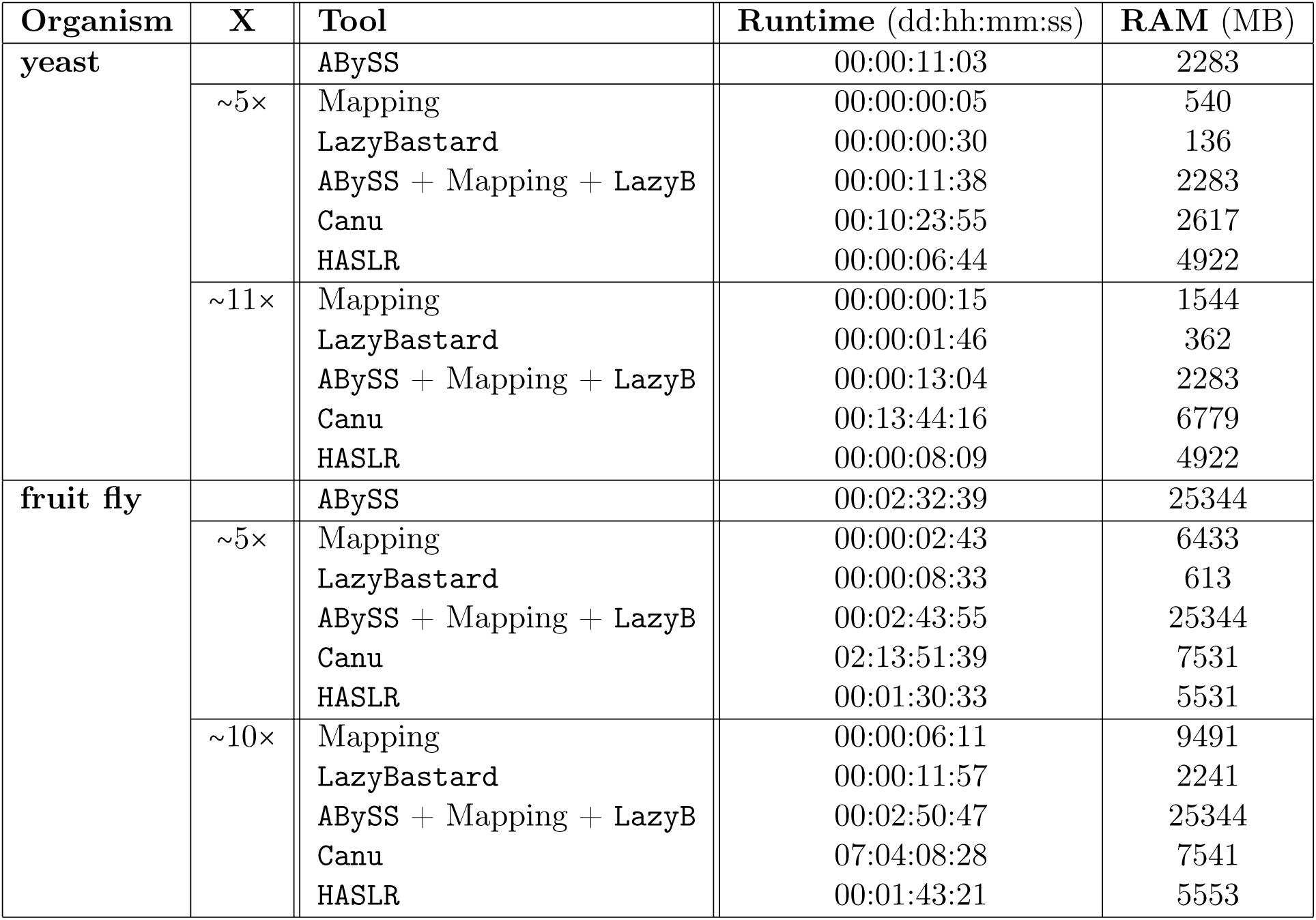
Assessment of running times for `LazyBastard` and `Canu`. Resources for `LazyBastard` are given in three steps: 1) `ABySS` unitig assembly; 2) Mapping of unitigs to long reads and 3) `LazyBastard` itself. Step 1) is often not needed as short-read assemblies are available for many organisms. Resources are only compared for yeast and fruit fly, as `Canu` cannot be run for human in sensible time and resource-constraint on our machine. As all tools except `LazyBastard` are parallelized, running times are given as the sum of time spent by all CPUs. `ABySS` greatly dominates the `LazyBastard` pipeline. Nevertheless, `LazyBastard` is faster on a factor of > 60.

The resource footprint of `LazyBastard` is small enough to run on an off-the-shelf desktop machine or even a laptop. The total effort for our approach is, in fact, already dominated by the computation of the initial unitig set from short reads. We expect that an optimized re-implementation of `LazyBastard` will render its resource consumption negligible. Compared to the competing `Canu` assembler, the combination of `ABySS` and the `python`-prototype of `LazyBastard` is already more than a factor of 60 faster. In terms of memory, `LazyBastard` also requires 3 − 18 less RAM than `Canu`; see Tbl. 2. Most notably, we were able to assemble the human genome within only 3 days, while `Canu` could not be run within our resource constraints. `HASLR` shows a similar distribution of running times between tasks, overall operating slightly faster.

## Discussion and Outlook

We demonstrated here the feasibility of a new strategy for sequence assembly with low coverage long-read data. Already the non-optimized prototype `LazyBastard`, written entirely in `python`, not only provides a significant improvement of the assembly but also requires much less time and memory than state-of-the-art tools. This is achieved by avoiding both a correction of long reads and an all-against-all comparison of the long reads. Instead we use rigorously filtered short-read unitigs as anchors to sparsifying the complexity of full string-graphs construction. Fast algorithms are then used to step-by-step orient the long reads and to remove inconsistencies from the already sparse graph, resulting in a set of paths corresponding to contigs.

The prototype implementation leaves several avenues for improvements. We have not attempted here to *polish* the sequence but only to provide a common coordinate system defined on the long reads into which the short-reads unitigs are unambiguously embedded to yield high-quality parts of the `LazyBastard`-assembly. The remaining intervals are determined solely by long-read data with their high error rate. Multiple edges in the multigraph constructed in the assembly step correspond to the same genome sequence, hence the corresponding fragments of reads can be aligned. This is also true for alternative paths between two nodes. This defines a collection of alignments distributed over the contig, similar to the situation in common polishing strategies based on the mapping of (more) short-read data or long reads to a preliminary assembly. Preliminary tests with off-the-shelf tools such as `racon` [23], however, indeed improve sequence identity but also tend to introduce new translocation breakpoints. We suspect this is the consequence of InDels being much more abundant than mismatches in Nanopore data, which is at odds with the NeedlemanâĂŞWunsch alignments used by polishing tools.

The mis-assemblies within the `LazyBastard` contigs were mostly inherited from chimeric reads. This therefore suggests an iterative approach: Subsampling the long-read set will produce more fragmented contigs, but statistically remove chimeric reads from the majority of replicate assemblies. Final contigs are constructed in a secondary assembly step by joining intermediary results. It might appear logical to simply run `LazyBastard` again to obtain a “consensus” assembly, where intermediary contigs play the role of longer reads with mapped anchors. In preliminary tests, however, we observed that this results in defects that depend on the sampling rate. The question of how to properly design the majority calling to construct a consensus assembly remains yet to be answered.

